# Total cerebral blood volume changes drive macroscopic cerebrospinal fluid flux in humans

**DOI:** 10.1101/2023.04.13.536674

**Authors:** Juliana Zimmermann, Clara Boudriot, Christiane Eipert, Gabriel Hoffmann, Rachel Nuttall, Viktor Neumaier, Moritz Bonhoeffer, Sebastian Schneider, Lena Schmitzer, Jan Kufer, Stephan Kaczmarz, Dennis M Hedderich, Andreas Ranft, Daniel Golkowski, Josef Priller, Claus Zimmer, Rüdiger Ilg, Gerhard Schneider, Christine Preibisch, Christian Sorg, Benedikt Zott

## Abstract

In the mammalian brain, the directed motion of cerebrospinal fluid (CSF-flux) is instrumental in the distribution and removal of solutes. Changes in total cerebral blood volume (CBV) have been hypothesized to drive CSF-flux. We tested this hypothesis in two multi-modal brain imaging experiments in healthy humans, in which we drove large changes in total CBV by neuronal burst-suppression under anesthesia, or by transient global vasodilation in a hypercapnic challenge. We developed a technique to monitor CBV changes based on associated changes in total brain volume by functional MRI (fMRI) and measured cerebral blood flow by arterial spin-labeling. Relating CBV-sensitive signals to fMRI-derived measures of macroscopic CSF flow across the basal cisternae, we demonstrate that increasing total CBV extrudes CSF from the skull and decreasing CBV allows its influx. Moreover, CSF largely stagnates when CBV is stable. Together, our results establish the direct coupling between total CBV changes and CSF-flux.

## Introduction

In the mammalian brain, the distribution and removal of substances depends on the exchange of molecules between the cerebrospinal fluid (CSF) and the interstitial fluid ^1–4^. In consequence, the directed movement of CSF (i.e., CSF-flux) into the brain parenchyma is necessary for the effective transport of solutes ^5^. CSF is formed in the choroid plexus, moves along the ventricles, basal cisternae, and subarachnoid spaces, where it enters the brain parenchyma via periarterial spaces^6^. However, we do not fully understand what drives CSF forward along this pathway.

Several drivers of CSF flux have been proposed, most prominently arterial pulsations and intra-cerebral pressure changes associated with the cardiac ^7–10^ and respiratory cycles ^11–14^. Additionally, accumulating evidence points towards infra-slow to slow cerebral hemodynamic changes as a third important driver of CSF-flux in the brain, especially in macroscopic compartments, leading to larger and slower CSF kinetics than those induced by arterial pulsations and respiration ^12,15^. Several functional magnetic resonance imaging (fMRI) studies in humans investigated the relationship between fMRI signals in the global grey matter (gGM) and in CSF-containing voxels of the fourth ventricle or basal cisternae ^16–19^, where, due to the so-called inflow effect, fluid entering the imaging volume appears brighter than stationary or exiting fluid ^17^. Recent evidence suggests that the hemodynamic changes associated with gGM fMRI signal fluctuations drive CSF motion and that such gGM hemodynamic changes can be induced by neuronal activity. Thus, slow coherent neuronal oscillations across the brain (i.e., delta waves of slow-wave sleep) are coupled to surges of CSF movement into the fourth ventricle ^17^. Moreover, visual stimulation, inducing a significant increase in neuronal activity and an associated hemodynamic change in the occipital cortex can drive macroscopic CSF flux across the basal cisternae ^15^. Given the Monroe-Kellie doctrine ^20^ stating that, under the condition of stable intracranial pressure within the rigid cavity of the skull, the volumes of blood, brain tissue and CSF must be constant, it is assumed that the observed gGM fMRI changes reflect total cerebral blood volume (CBV) alterations, which lead to anticorrelated cranial in- or outflow of CSF ^17,21^.

So far, however, direct evidence that slow changes in total CBV drive CSF-flux is missing, because direct experimental manipulation and time-resolved measurement of total CBV is challenging. Thus, current evidence is based on indirect manipulations of CBV via neuronal activity ^15,17^ or forced inspiration, putatively altering the intracranial pressure ^16^, but in both cases direct evidence for CBV, particularly total CBV changes is missing. Instead, previous studies mainly relied on the gGM fMRI signal as a proxy for total CBV. This may be problematic as fMRI signal changes in the grey matter are primarily linked to changes in the level of blood oxygenation rather than CBV ^22,23^, potentially confounding the assumed blood volume relationships with CSF-flow signals.

To address these issues, we performed two related experiments to investigate the effects of total CBV changes on macroscopic CSF flux across the basal cisternae by combining experimental conditions, which ensure maximal total CBV changes, with direct and time-resolved total blood volume-sensitive measurements. Concretely, we first used simultaneous EEG-fMRI during deep anesthesia of burst-suppression in healthy young volunteers. The burst-suppression-EEG pattern is suited to study the effects of total CBV on CSF flow because it constitutes instantaneous, brain-wide contrasts between isoelectric suppression phases (i.e., minimal neuronal activity) and global bursts (i.e., high broadband-power activity) (Shanker et al 2021). This study expands previous work during slow-wave sleep (Fultz et al 2019, Picchioni et al 2022) or wakefulness (Williams et al 2023, Yang et al 2022), in which global brain activity is partly and slowly changed with respect to its spatial-temporal activity pattern (Cash et al 2009). From the fMRI, we extracted the CSF signal at the bottom slice of the imaging volume as well as the gGM signal. Additionally, based on the observation that brain-wide changes in CBV are associated with measurable alterations of total brain (i.e., parenchyma) volumes ^24,25^, we developed a new technique to track CBV-induced changes in total brain volume. To this end, we analyzed the fMRI signal time course at the interface between the brain parenchyma and the ventricular CSF to detect shifts of the parenchyma-fluid interface (PFI) induced by burst-suppression associated changes in total brain volume.

Second, we confirm and expand the findings in a further experiment in which we directly manipulated total CBV by performing a controlled hypercapnic challenge, i.e., transiently elevated inspiratory CO2 levels. Hypercapnic challenges cause global vasodilation and thus global increases in cerebral blood flow (CBF) (i.e., brain perfusion) and total CBV in animals ^26^ and humans ^27^ at largely stable or even slightly decreased global neuronal and metabolic activity ^28,29^. To confirm the vasodynamic effects of the hypercapnic challenge, we performed time-resolved pseudo-continuous arterial spin-labeling (pCASL) MRI to extract the CBF time courses, which we use as an indicator for concurrent CBV changes due to the well-established relationship between the two ^30–32^. Then, under the same experimental conditions of confirmed total CBV changes and assumed stable oxygen metabolic neural activity, we used blood volume sensitive fMRI and observed large transient changes in the gGM fMRI signal tightly coupled to flow-dependent CSF fMRI signal alterations.

Finally, consolidating both experiments, we define transition events (burst Η suppression, hypercapnia Η normocapnia) as well as steady-state phases (burst or suppression, hypercapnia or normocapnia) to study the consequences of induced gGM-CSF-coupling for CSF-flux into and out of the brain under such different conditions of global brain states (e.g., spontaneous breathing in wakefulness and mechanically ventilated anesthesia) or different time scales (e.g., burst-suppression transitions of milliseconds and hyper-normo-capnia transitions of 30 seconds).

## Results

### Globally coherent neural activity switches drive CSF-flux, mediated by CBV

First, we tested the effects of globally coherent neuronal activity switches on gGM, PFI and CSF fMRI signals and their coupling under burst-suppression anesthesia. We used previously recorded data from intubated and deeply sevoflurane-anesthetized (∼4,3%) healthy subjects (n = 17) ^33^ under a burst-suppression EEG pattern ^33–35^. EEG and fMRI data were co-registered in a 3T-MRI scanner (**Fig. 1a**).

**Fig. 1.**
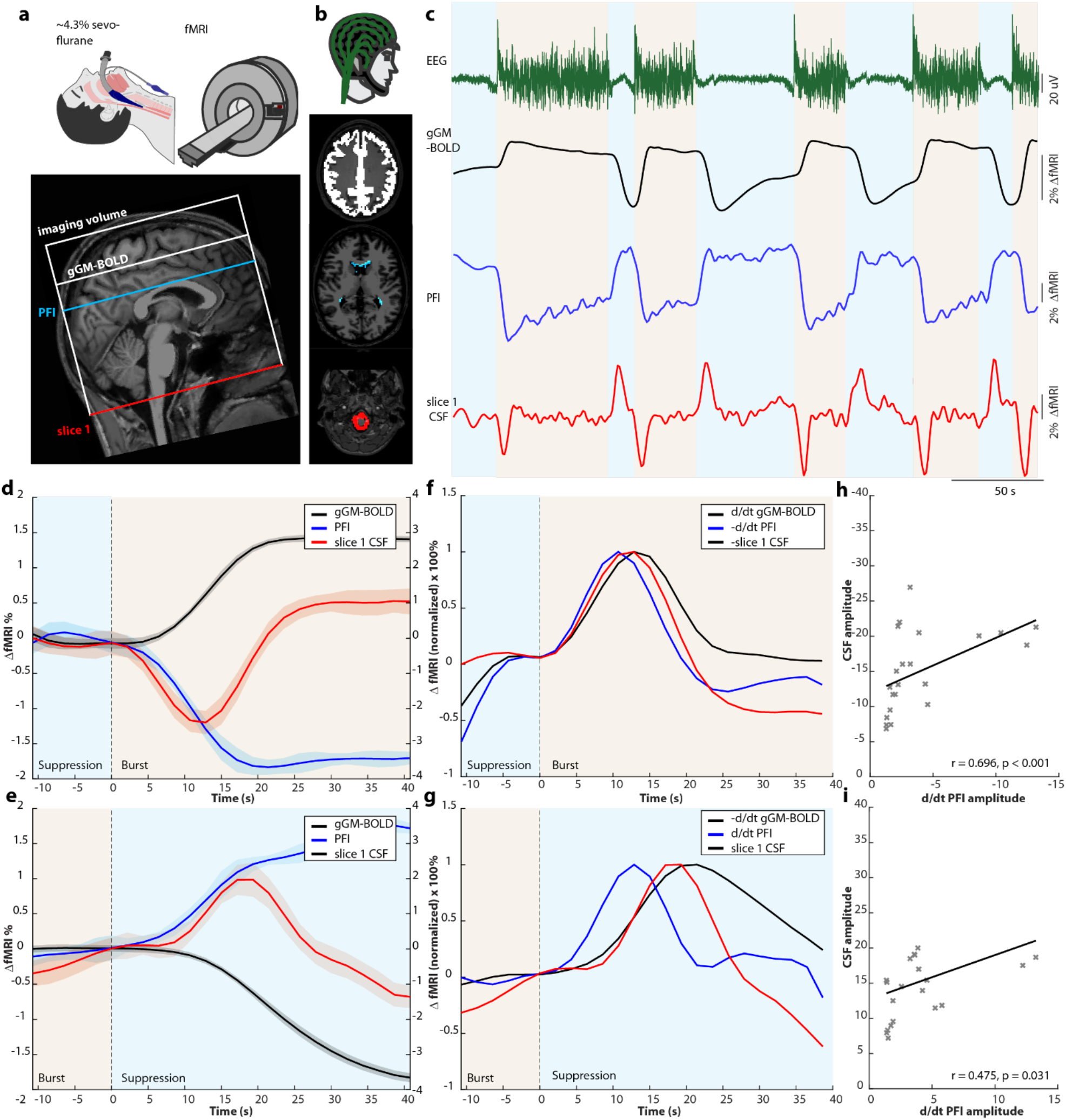
Coupled neuronal activity, gGM, PFI and CSF fMRI signals during burst-suppression anesthesia. (**a**) *top*: Experimental design: fMRI recording under deep sevoflurane anesthesia. Bottom: Example scan positioning in a representative subject (#5). Schematic depiction of the fMRI volume superimposed on a sagittal T1-weighted image with representative slice positions for the gGM and PFI masks, as well as slice 1, which contains the CSF mask. (**b**) Simultaneous EEG and fMRI recordings of three masks, depicted in representative slices: gGM (*white*), PFI (*blue*) and CSF (*red*). (**c**) Simultaneously recorded signal time courses from one subject under burst-suppression anesthesia: EEG (*green*), gGM fMRI (*black*), PFI fMRI (*blue*) and CSF fMRI (*red*). Suppression and burst periods are indicated in light blue and light orange, respectively. (**d**) Subject- and event-averaged time courses of all transitions from suppression to burst (n=22 events from 17 subjects). gGM fMRI (*black*), PFI fMRI (*blue*) and CSF fMRI (*red*). The shaded areas represent the standard error of the mean (SEM). Suppression and burst periods are indicated in light blue and light orange, respectively. The y-axis on the right contains y-values for the PFI and CSF fMRI signal. (**e**) Same as (d) for transitions from burst to suppression (n = 21). (**f**) Event-averaged time courses of d/dt gGM fMRI, -d/dt PFI fMRI and -CSF fMRI, normalized to 100% for suppression-burst events. (**g**) Event-averaged time courses of -d/dt gGM fMRI, d/dt PFI fMRI and CSF fMRI, normalized to 100% for burst-suppression events. (**h**) Scatterplot of the d/dt PFI and the CSF amplitude. Crosses represent transition events; the line depicts linear regression. (**i**) Same as (h) for all burst-suppression events.

From the echo planar imaging (EPI) fMRI signal of each individual, we generated three masks (**Fig. 1b**). First, we defined a gGM mask to extract the fMRI signal as a marker for global blood-oxygen level-dependent and hemodynamic-volumetric changes in the brain ^15,17^. Second, we defined the mask of the PFI to obtain a marker for changes in total brain volume as a function of CBV, which, unlike the gGM fMRI signal, is not affected by blood oxygenation levels. We generated the PFI mask in each subject individually by subtracting the raw image of the entire volume stack averaged across all suppression epochs from the volume averaged across burst epochs (**Fig. S1a-d, Methods**). This resulted in a ring-shaped pattern of voxels with negative intensity values at the borders of the lateral ventricles (**Fig. S1d-g**). Due to the high contrast in echo planar imaging signals of fMRI between CSF (high intensities) and brain parenchyma (low intensities) (**Fig. S1b, c**), this indicates that during bursts, this ring of voxels contains a higher proportion of brain parenchyma than during suppression, causing a partial volume effect. We conclude that, during bursts, the brain parenchyma expands, thus slightly compressing the lateral ventricles (**Fig. S1h**). In line with this, a ring of voxels with negative subtraction values was also detectable over the convexity of the skull (**Fig. S1i**), indicating that the expanding brain parenchyma during bursts displaces both ventricular and subarachnoid CSF. Third, we defined the CSF voxels of slice 1, the bottom slice of our imaging volume, containing the premedullary, lateral and posterior cerebellomedullary cisternae (**Fig. 1b**) to detect inflow-related CSF signal changes^15,17,19^.

From all subjects, we extracted the EEG time course as well as the simultaneously recorded fMRI signals from our three defined masks. On the subject level, we observed changes in all three signal time courses, which were time-locked, with a slight delay, to each individual suppression-burst and burst-suppression transition in the EEG (**Fig. 1c, Fig. S2 for an additional subject**). Thus, the gGM fMRI signal steeply increased with the onset of each burst and rapidly decreased after the switch to suppression, in line with previous reports ^35^. The PFI signal showed a bi-stable signal time-course with low signal intensities during bursts and high intensities during suppression. A control analysis confirmed that the PFI signal was not affected by signals in the neighboring CSF or the white matter (**Fig. S3**), reinforcing that it is indeed driven by rapid changes in parenchyma volume. In the CSF signal in slice 1, we detected a positive peak associated with burst-suppression switches and a negative peak after suppression-burst switches. The observed CSF signals in slice 1 were not influenced by blood flow through the vertebral arteries in the region of interest (**Fig. S4).** In contrast to the rapidly changing fMRI signals at neuronal activity transitions, the CSF and gGM fMRI signals were comparatively stable during phases of unchanging neuronal activity, i.e., neuronal suppression epochs or long bursts (**Fig. S5**).

Next, we averaged all three fMRI signals from the burst-suppression (n = 21 events) and suppression-burst (n = 22) transitions (**Fig. 1d, e, methods**) in all subjects. This confirmed the strong association between the three signals already visible at the single event level (**Fig 1 c**), i.e., in the gGM and PFI fMRI signals, an anti-correlated bi-stable signal time course with relatively rapid state-switches following the neuronal transitions with a short delay. The CSF signal showed a negative dip following suppression-burst transitions with a maximum amplitude 12.9 s after the transition and a positive peak following the bust-suppression transition after 17.2 s.

To analyze temporal dynamics between PFI, gGM and CSF fMRI signals, we calculated d/dt of the PFI and -d/dt of the gGM fMRI signal ^17^. The -d/dt PFI signal preceded the CSF signal under both conditions with a longer lag for burst-suppression compared to suppression-burst transitions (**Fig. 1f, g**). On the other hand, there was no obvious delay between the gGM and the CSF fMRI signal, possibly due to the long TR in our recordings. Cross-correlation analysis revealed that the PFI signal preceded both the gGM and the CSF fMRI signals on subject- and single-event levels and confirmed that there was no delay between the gGM and CSF fMRI signals (**Fig. S6**).

If the gGM and/or the PFI fMRI signal were to drive CSF-flux, one would expect that large PFI changes would drive large CSF flow events. To test this, we correlated the amplitude of the CSF with the d/dt PFI and d/dt gGM fMRI signals. We found a correlation between the amplitudes of the d/dt PFI and CSF signals, both for suppression-burst (**Fig. 1h**) and burst-suppression **(Fig. 1i**) transitions. In contrast, there was no association between d/dt gGM and CSF fMRI signal amplitudes (**Fig. S7**). Together, these data indicate that, at neuronally induced burst-suppression transitions, PFI and the associated change in CBV drive macroscopic CSF flow in the basal cisternae.

### Hypercapnia-induced vasodilation drives CSF flow via changes in cerebral blood flow

In a separate experiment, we investigated whether the experimental manipulation of vascular diameter can drive CSF motion. We utilized a controlled hypercapnic challenge design in healthy subjects (n=17) to induce transient cerebral vasodilation under awake resting-state conditions ^36^. Within a total scan duration of 900 sec, baseline periods (180 s) alternated with hypercapnia periods (5% inspiratory CO2). The transitions between both states took ∼30 s. (**Fig. 2a**). CBF and fMRI signal changes were acquired separately by optimized pCASL and fMRI sequences (**methods**).

**Fig. 2.**
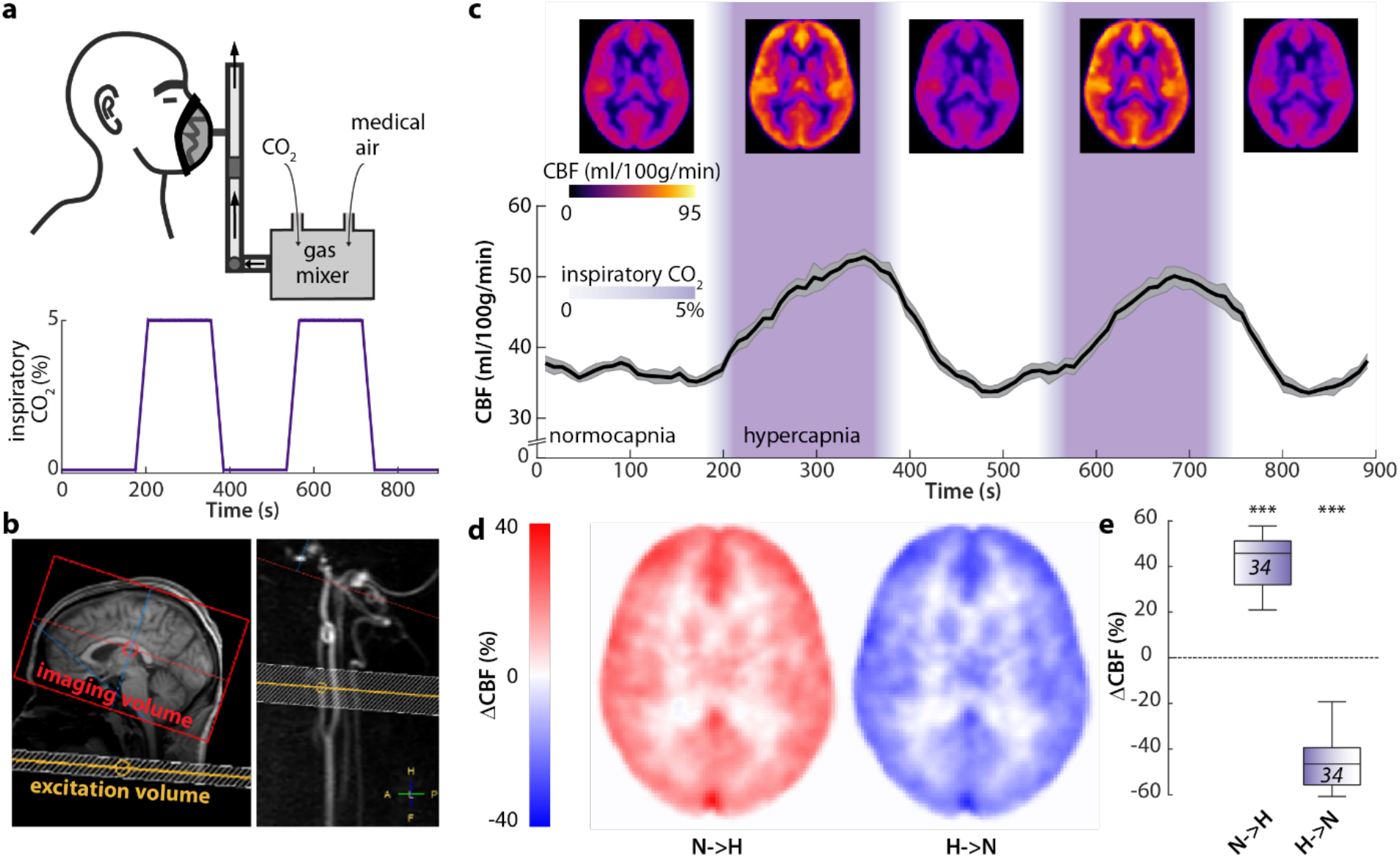
Bidirectional grey-matter CBF changes induced by a hypercapnic challenge. (**a**) Schematic depiction of the experimental design: a gas mixture of CO2 and medical air was applied over a medical mask in awake subjects *(top)*. Schematic time course of the inspiratory CO2 concentrations of the applied gas mixture including two 180 s segments with elevated CO2 levels (5% Vol/vol). Transition periods lasted ∼30s due to ramp time of the gas mixer. (**b**) Example scan positioning for the pCASL imaging volume (red) and labeling plane (yellow) in a representative subject superimposed on a sagittal T1 image (*left)* and a sagittal reconstruction of a 3D-phase contrast angiogram (*right*). (**c**) Whole brain cerebral blood flow (CBF) maps (*top*) and extracted subject-average (n = 17) time course of gGM CBF (*bottom*). Mean (*black*) ± SEM (*grey*). CO2 application periods are color-coded in *purple*. (**d**) Subject-average maps of CBF changes for transitions from normocapnia to hypercapnia *(N→H, left)* or from hypercapnia to normocapnia (H→N, *right*). (**e**) Global grey matter ΔCBF values for *N→H (n =34, left)* or *H→N (n =34, right) transitions* (2 for each subject). One-sample t-test (individual groups). ***p<0.001.

In a first step, we evaluated pCASL MRI data for CBF to investigate the effect of the hypercapnic challenge due to simultaneous vasodilation/vasoconstriction and perfusion increase/decrease (**Fig. 2b**). Total CBF was high across the whole brain during vasodilation periods of 5% CO2 compared to zero CO2 conditions (**Fig. 2c, *top***), which is in line with previous studies ^36–38^. Time-resolved analysis of the global grey matter CBF (gGM-CBF) showed that it reliably followed the levels of inspiratory CO2, i.e., was relatively stable during baseline conditions, increased with the rise of CO2, until it almost reached a plateau, before decreasing with falling CO2 (**Fig. 2c, *bottom,* Movie S2**). The gGM-CBF increased by 41.6 +/- 11.8% from normo- to hypercapnia and decreased by 44.8 +/- 13% during the inverse transition (**Fig. 2 d,e, Movies S3, S4**). Critically, due to hypercapnia-induced vasodilation together with massive total perfusion increase and decrease, we can infer total CBV changes during the hypercapnic challenge.

Second, we evaluated the fMRI data for gGM and cisternal CSF fMRI signals and their coupling in the same subjects during an identical hypercapnic challenge (**Fig. 3a**). We extracted fMRI signals from gGM as well as from CSF-containing voxels of slice 1. Note that the hypercapnic challenge leads to GM CBV increase under rather stable or even reduced metabolic neuronal activity, suggesting the gGM fMRI signal to reflect mainly blood volume changes. On single subject (**Fig. S8**) and group levels (**Fig. 3b**), upon the onset of hypercapnia, the gGM fMRI signal increased from a stable baseline and almost reached a plateau during the time at which CO2 was kept at 5%, before decreasing again following the drop of inspiratory CO2 levels. In the CSF-containing voxels of slice 1, we detected a signal dip that co-occurred with the increase of the gGM fMRI signal (**Fig. 3c)** and a positive peak associated with the decrease of gGM fMRI signal (**Fig. 3d**). The gGM and cisternal CSF signals were anticorrelated during both transition periods from normo- to hypercapnia (N→H) and from hyper- to normocapnia (H→N) (**Fig. 3e**).

**Fig. 3.**
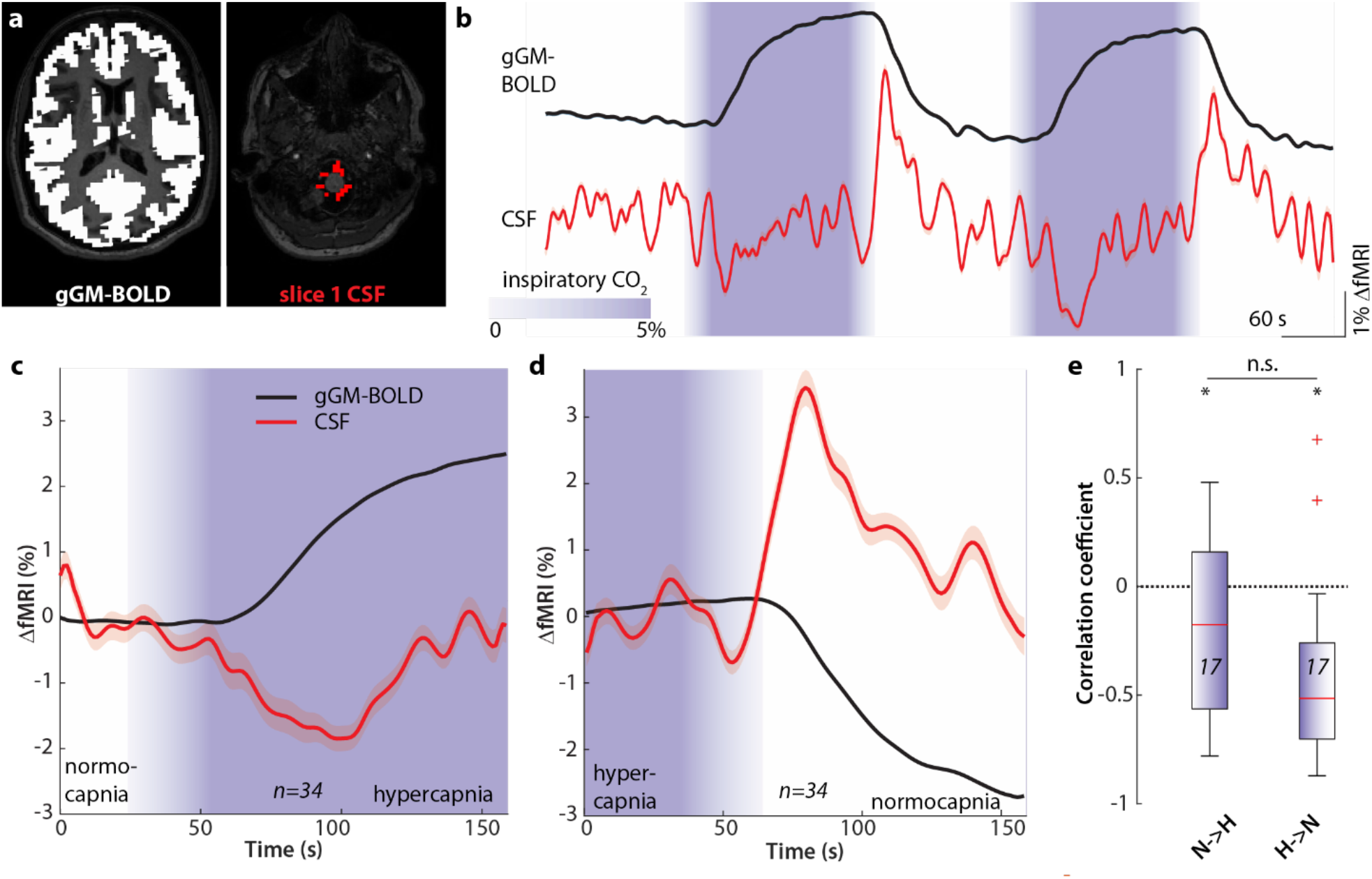
Coupled gGM and CSF fMRI signal changes induced by a transient hypercapnia challenge. (**a**) Global grey matter (*left*) and CSF masks *(right)* for extracting fMRI signals, superimposed on axial T1-weighted images. (**b**) Time courses of the gGM *(top, black)* and CSF *(bottom, red)* fMRI signals. Mean (*solid lines*) ± SEM (*shaded areas*) of n = 17 subjects. CO2 application periods are color-coded in purple. (**c**) Averaged time courses of all normocapnia to hypercapnia (N→H) transitions (n = 34 transitions from 17 subjects). gGM *(black)* and CSF fMRI signal *(red)*. The shaded areas represent SEM. CO2 application periods are color-coded in purple. (**d**) Same as (C) for transitions from hypercapnia to normocapnia (H→N, n = 34 transitions from 17 subjects). (**e**) Correlation coefficients between gGM and CSF fMRI signals during N→H *(n=17, left)* and H→N *(n=17, right)* transitions. *p<0.05, n.s. not significant. One-sample t-test (individual groups). Two-sample t-test (between groups).

In summary, these results demonstrate that H→N and N→H transitions, respectively, induce total CBV changes, which underpin gGM fMRI signal changes coupling inversely with CSF signal changes in the basal cisternae of the brainstem.

### Macroscopic CSF in- and outflux due to CBV changes

Finally, across both experiments of burst-suppression during anesthesia and transient hypercapnia during wakefulness, we analyzed the CSF signals with respect to indicators of flux direction using slice-sensitive fMRI analysis of fMRI-based CSF signals in the basal cisternae (**Fig. 4a**). In both experiments, we detected a transient increase in CSF signal intensity that co-occurred with the gGM fMRI signal drop (**Figs. 4 b,c**). In line with the previous study of Fultz et al. ^17^, the CSF signal amplitude decreased with increasing slice number in both experiments (**Fig. 4 b,c**), more steeply in the burst-suppression experiment. This amplitude decrease is indicative of influx into the imaging volume, as water-protons (‘spins’), which are unsaturated and hence hyperintense when entering the imaging volume, get increasingly saturated and thus less hyperintense by experiencing repeated radiofrequency pulses along slices above the bottom slice ^39^.

**Fig. 4.**
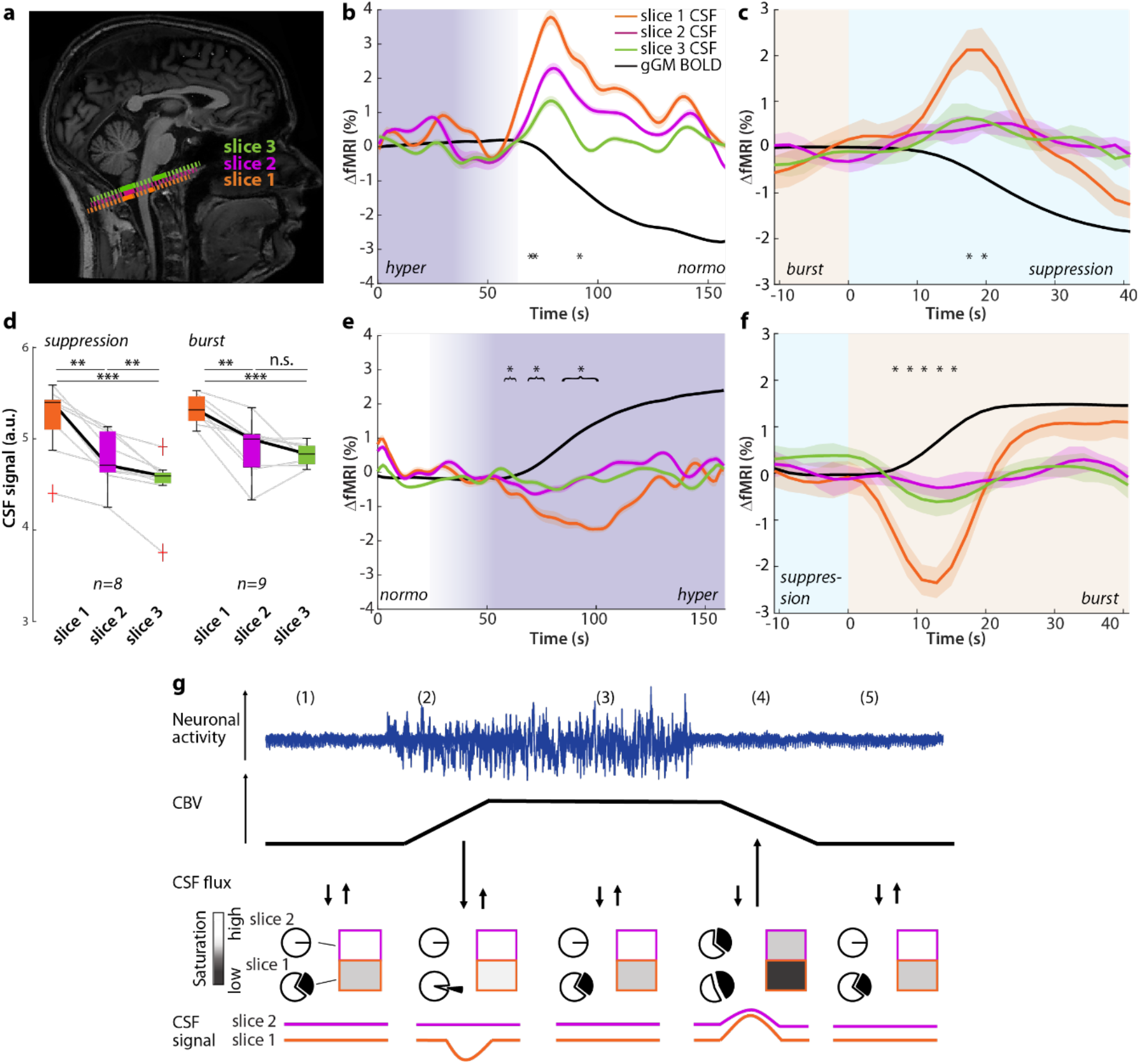
Changes in global brain blood volume mediate CSF in- and outflux. **(a)** Representative example of the positioning of the most caudal fMRI slices on a sagittal T1-weighted image. The CSF voxels of slice 1 are indicated in *orange,* slice 2 in *pink* and slice 3 in *green,* the dashed lines are schematic depictions of the respective imaging slices. (**b)** Average time courses of all transitions from hypercapnia to normocapnia. gGM (*black*), CSF fMRI signal in slice 1 (*orange*), slice 2 (*pink*) and slice 3 (*green*). The shaded areas represent SEM. Hypercapnia and normocapnia periods are indicated in purple. *p<0.05, repeated-measures ANOVA (between slices) per subject (n=17). **(c)** Average time courses of all transitions from burst to suppression. Global GM (*black*), CSF fMRI signal in slice 1 (*orange*), slice 2 (*pink*) and slice 3 (*green*). The shaded areas represent SEM. Burst and suppression periods are indicated in light orange and light blue, respectively. *p<0.05, repeated-measures ANOVA (between slices) per event (n=21). (**d)** Mean fMRI-based CSF signal intensities (a.u.) per event of slices 1-3 for steady-state suppression or steady state burst periods. ** p<0.005, ***p<0.001, n.s. not significant. Repeated measures ANOVA with Dunn-Sidak post-hoc comparison. (**e)** Same as (b) for all transitions from normocapnia to hypercapnia. *p<0.05, repeated-measures ANOVA (between slices) per subject (n=17). (**f)** Same as (C) for all transitions from suppression to burst. ** p<0.005, ***p<0.001, repeated-measures ANOVA (between slices) per event (n=22). **(g)** Schematic model of the link between neuronal activity (*blue*), CBV (*black*), CSF flux (↓ - efflux from the acquisition volume, ↑ - influx into the acquisition volume), fMRI-based CSF signal in slice 1 (*orange*) and slice 2 (*pink*). The pie charts symbolize the saturation state of CSF, being either fully saturated (*white*) or unsaturated (*black*); the rectangles indicate the signal intensity in the respective slice, which results from the mixing of RF-saturated and fresh CSF.

During the transition from suppression to burst or from normo- to hypercapnia, the transient signal dip in the fMRI-based CSF signal is associated with an increase in gGM fMRI signal that is clearly detectable in slice 1 but largely attenuated in slices 2 and 3 in both experiments (**Fig. 4 e,f**). This signal dip likely reflects the passage of attenuated CSF through slice 1 ^39^, as would be expected by the extrusion of saturated CSF from the imaging volume. Accordingly, during steady-state conditions in the first experiment, i.e., phases of quiescent suppression or during long bursts, the analysis across several slices revealed that the CSF signal intensity of slice 1 was higher than that of slices 2 and 3, for both long bursts and long suppression phases (Fig. 4d**, Fig. S9**).

To further investigate whether the observed bidirectional changes in fMRI-based CSF signal intensity were associated with in- and outflux of CSF, we compared the traces of slice 1 CSF signal to the negative derivative of the gGM fMRI signal (**Fig. S10, A-F**). In previous reports ^17,21^, the CSF signal closely resembled the -d/dt gGM fMRI signal, in which negative values were set to zero, indicating that only influx of CSF could be detected. However, our analysis revealed that for both experiments, the correlation between the slice 1 CSF signal and the non-thresholded -d/dt gGM fMRI signal was higher than the zero-thresholded version, indicating that the signals were caused by in-and outflux rather than only by influx. Interestingly, comparing both datasets, we found that the derivatives, i.e., slopes, of the leading flank of the CSF and the gGM fMRI signals were steeper for the burst-suppression data, compared to the hypercapnia data (**Fig. S10G**), confirming the observed faster transitions in the former.

Together, these results provide strong evidence for the notion that total CBV changes associated with global neuronal activity or experimentally induced by a hypercapnic challenge cause bidirectional in- and outflux of CSF across the basal cisternae.

## Discussion

This study demonstrates, firstly, that rapid switches between cortical quiescence and phases with bursts of strong global neuronal activity during deep anesthesia induce large global changes in the total brain volume, reflective of CBV fluctuations, which cause inversely related macroscopic in- or outflux of CSF across the basal cisternae with a slight delay. Secondly, we provide evidence from a hypercapnic challenge experiment during wakefulness that gGM fMRI and CBF signal changes during hyper- to normocapnia or normo- to hypercapnia transitions drive bidirectional CSF motion into and out of the skull. Since such vasodilation-associated CBF changes go along with CBV changes ^36–38^, our results indicate that global CBV is a direct driver of CSF flux.

Integrating these observations leads us to a unified model of global neuronal activity changes and related brain CBV-induced flux of CSF into and out of the brain (**Fig. 4g**), which both confirms and expands previous models based on CSF influx data only ^17,21^. To explain this model, we start with steady-state baseline conditions – such as ongoing suppression (**Fig. 4g, *1***). During suppression, there are no large surges of CSF across the basal cisternae (**Fig. S1**). However, the CSF signal intensity in the lowest slice 1 is slightly higher than that of slice 2 (**Fig. S9**), likely due to a constant exchange with ‘fresh’ less saturated CSF at the edge of the imaging volume. This could be due to small inflow events caused, among others, by respiration and heartbeat ^12,13^, which reached the lowest, but not the adjacent slices, and could not be resolved due to lack of temporal resolution in our imaging paradigm. When global neuronal activity increases - as in the transition from suppression to burst (**Fig. 4g, *2***) -, this rise causes an increase in CBV (**Fig. 2**). The accompanying increase in brain volume extrudes less saturated CSF spins from slice 1 and replaces them by more saturated spins from fluid inside the imaging volume, thus leading to a transient signal dip in slice 1 but not in more cranial slices.

It is important to note that a negative signal transient in slice 1, which, according to our model, represents outflux, was not reported previously ^16–19,21,40^. However, these studies used experimental conditions with less pronounced changes of gGM-BOLD signal or patients with severe brain disease. During the burst, CBV remains elevated (**Fig. 4g, *3***) and comparatively stable. Accordingly, steady-state conditions are established in the CSF, where again, fMRI signal intensity in slice 1 is higher than in slice 2. The instantaneous switch to neuronal suppression (**Fig. 4g, *4***) causes a decrease in CBV (and brain volume), leading to a transient inflow of fresh unsaturated fluid from outside the imaging volume into the skull. This leads to a pronounced inflow signal with a high amplitude in slice 1, which is largely attenuated in higher slices (**Fig. 4**), most likely, because of saturation of the entering CSF spins by experiencing an increasing number of RF-pulses ^17^. After CBV has returned to baseline levels (**Fig. 4g, *5***), a steady state is reached again.

Beyond this model, the observed coupling of CBV changes to CSF in- and outflux is comparable in both experiments despite clear differences in methods, context, and dynamics. First, CBV-CSF coupling was induced either by changes in neuronal activity during sevoflurane anesthesia or, in awake subjects, by vasodilation induced by the hypercapnic challenge with largely stable or even slightly decreased neuronal activity ^28,29^. Second, under sevoflurane anesthesia, the participants were ventilated mechanically by positive inspiratory pressure instead of negative pressure upon spontaneous breathing in the hypercapnia dataset. Third, sevoflurane anesthesia causes general cerebral vasodilation ^41^, which may dampen the amplitude of the response to neuronal activation. Third, the experiments induced CSF flow at different timescales. Thus, the rapid CBV changes associated with instantaneous neuronal burst-suppression transitions drive short CSF signal peaks and dips with high signal slopes indicative of fast CSF flow ^39^, while the slow vasodilation-driven CBV changes in the hypercapnia dataset also drove more extended CSF peaks with lower slopes. Despite these differences, both experiments showed similar coupling between our CBV proxy measures with CSF flow. This indicates that the described mechanism of CBV-CSF coupling is a fundamental physiological principle which is robust across very different conditions.

Linking results with brain waste clearance or more generally with metabolite distribution, bulk motion of CSF in the basal cisternae is tightly coupled to that in the 4^th^ ventricle ^18^ and may be related to an increase in perivascular flow, which could facilitate brain waste clearance due to enhancing mixing and ‘directed diffusion’ or convective flow ^2^. In fact, recent evidence indicates that visual stimulation, which induces macroscopic CSF motions in the 4^th^ ventricle in humans ^15^, is associated with an increased flow velocity in perivascular spaces in rodents ^9^. If so, burst-suppression transitions may via their macroscopic CSF flux effects exert a protective clearance function in severe brain pathologies such as early onset infantile epilepsy, coma or hypothermia^42^. Interestingly, medically-induced burst-suppression is used as a neuroprotective tool in severe epilepsy ^43^. However, under surgical anesthesia, burst-suppression states are not desired and associated with detrimental outcomes ^42^.

Under physiological conditions, similar yet less drastic fluctuations in CBV or gGM fMRI signal can be detected in slow-wave sleep ^17^, awake resting-state conditions ^19^ or experimentally induced by visual stimulation ^15^. Remarkably, these fluctuations are also coupled to CSF movement across the basal cisternae or the 4^th^ ventricle, indicating that the coupling mechanism relies on a general physiological principle. Pathologically, uncoupling of CSF and CBV may contribute to neurodegenerative diseases including Alzheimer’s, possibly leading to a buildup of β-amyloid ^44^. The disruption of sleep slow waves and the associated clearing mechanisms ^1,45^ in Alzheimer’s ^46,47^ may further exacerbate this process.

The main limitation of our study is that we cannot entirely exclude that, in addition to the CBV changes, the CSF flow signal is influenced by other factors of systemic physiology, including heartbeat and respiration, which may arise simultaneously with the bursts-suppression or normo-hypercapnia transitions in our experiments. In fact, the relationship between CSF flow and cardiac pulsations and respiration is firmly established to lead to undulating movements of CSF ^7,8,10,13,14^, with faster kinetics than those observed here ^15^. However, an influence of changes in respiratory patterns can be excluded, because in our burst-suppression experiment, the subjects were mechanically ventilated at fixed respiration rates and tidal volumes ^33^. While mechanical ventilation uses positive inspiratory pressure, which may affect intracranial pressure and CSF flow ^48^, one would expect any effects of this to be constant during the entire imaging period. In contrast, the CSF signals were phase-locked to spontaneous burst-suppression transitions, which occurred at no apparent pattern, and lasted several to tens of seconds, much longer than respiration-induced CSF inflow signals ^15^. Second, while we did not directly measure the heart rate in our subjects, hypercapnic challenges did not cause systematic changes in heart rate in previous studies ^27,49^, excluding it as a key confounder for CSF flow in this dataset.

A second limitation is that the PFI signal is, in fact, a dynamic measure of total brain volume, rather than CBV only. However, while we cannot formally exclude that a small proportion of the PFI signal is caused by volume changes of structures other than blood vessels, including the interstitial or cellular compartments, the rapidity of the state-switches at burst-suppression transitions makes this very unlikely. Finally, in the hypercapnia dataset, we qualitatively estimate changes in CBV from dynamic CBF and gGM fMRI measures. Multiple studies have reported that in hypercapnic challenges, CBV closely tracks CBF ^30–32^. Moreover, hypercapnic challenges are associated with largely stable or even slightly decreased brain metabolism ^28,29^, indicating that under these particular conditions, the fMRI-signal is a better marker of hemodynamics than under the typical task-fMRI paradigms which strongly affect oxygen metabolism ^22^. This notion is also supported by the parallelism of the CBF and gGM fMRI signals in our experiment.

In summary, our experiments provide direct evidence that changes in total brain blood volume drives macroscopic CSF flux in healthy human subjects under different conditions.

## Methods

### Experimental Design

Data from two independent studies in healthy adults were used for the current study. Both were conducted in line with the Declaration of Helsinki and approved by the ethics committee of the medical school of the Technical University of Munich. Participants were given detailed information about the methods and potential risks and gave their written informed consent before the experiments.

### Dataset 1: Anesthesia experiment

#### Participants and anesthesia procedure

Data were derived from a previously published simultaneous EEG-fMRI study about sevoflurane effects on brain activity in healthy adults conducted at the Technical University of Munich, Germany ^33–35^. A detailed description of the participants and study protocol is reported in ^33^. In brief, combined EEG-fMRI measurements under sevoflurane-induced anesthesia were carried out in 20 healthy adult males from 20 to 36 years (mean 26 years). Data from 17 subjects is included in this study; data from three subjects was excluded because of missing fMRI acquisition markers (i.e., triggers), the absence of a clear burst suppression pattern in the EEG, or due to inadequate positioning of the imaging volume. Sevoflurane anesthesia was administered in oxygen via a facemask using MRI-compatible anesthesia monitoring equipment (Fabius Tiro, Dräger, Germany). Sevoflurane, as well as O2 and CO2 concentrations, were measured by a cardiorespiratory monitor (Datex AS/3, General Electric, USA). Sevoflurane was increased from 0.4% to 3%, and mechanical ventilation via a laryngeal mask (i-gel, Intersurgical, United Kingdom) was initiated when clinically indicated. Sevoflurane concentration was gradually increased. When the EEG showed a burst-suppression pattern with suppression periods of at least 1 second and about 50% suppression rate (4.34 +/-0.22 vol%), EEG-fMRI recordings of burst-suppression were performed.

#### EEG data acquisition and pre-processing

EEG was recorded using an fMRI-compatible 64-electrode cap (Easycap, Germany) and two 32-channel amplifiers (BrainAmp MR, Brain Products, Germany). Electrode impedance was kept below 5kΩ using an abrasive gel (Easycap). All signals were recorded at 5 kHz sampling rate. An interface unit (SyncBox, Brain Products) was additionally connected to the amplifiers to reduce timing-related errors in the fMRI artifact correction by synchronizing the clocks of the EEG amplifiers and the fMRI gradients. One of the 64 channels was placed over the chest for electrocardiography (left anterior axillary line).

EEG data preprocessing (BrainVision Analyzer 2.2.1) included automatic gradient artifact correction (MR correction) using a template drift detection method (TDC), low-pass FIR filtering to 40 Hz, and down-sampling by a factor of 20.

#### Outcome: Burst and suppression definition

The periods of burst and suppression used in fMRI data evaluation were defined based on the simultaneously recorded EEG traces described in (*24*).

Briefly, a semi-automatic labeling approach was used to detect burst and suppression episodes using one EEG channel: *(i)* visual assessment and confirmation of the presence of burst suppression pattern (i.e., alternating high-voltage activity with isoelectricity); *(ii)* FFT signal linear filtering (1-5Hz) using EEGLAB v2022.1; *(iii)* signal amplitude normalization (z-score); *(iv)* calculation of the averaged upper envelope of the sequence using the Matlab (R2020b, MathWorks, Natick, MA, USA) function ‘envelope’; *(v)* visual inspection and labeling of fMRI volumes with a 1 (i.e., burst) if the EEG signal amplitude surpassed at least two standard deviations above the mean in at least half of the entire 2s repetition time (TR; of fMRI volume), everything below was labeled with a 0 (i.e., suppression).

Based on the digital labels from the EEG traces, we extracted four types of events from the fMRI data of individual subjects: (i) epochs of suppression, (ii) epochs of burst, (iii) transitions from suppression to burst (S→B), (iv) transitions from burst to suppression (B→S). To ensure that the fMRI signals reached steady-state conditions before and after the respective events, we analyzed only B→S or S→B transitions, which occurred at least 20 s after the preceding and at least 40 s before the following transition. Periods of steady state burst‘ and ‘steady state suppression‘ were defined as continuous periods of >120 s (burst) and 160s (suppression), respectively. We omitted the first 40 s (burst) or 20 s (suppression) to allow for reaching a steady state.

#### MRI Acquisition and Preprocessing

Data was acquired in a 3T whole-body MRI scanner (Achieva Quasar Dual 3.0T 16CH; Philips, Medical Systems International Inc., Best, Netherlands) with an eight-channel, phased-array head coil. First, a T1-weighted magnetization-prepared rapid gradient echo (MPRAGE; voxel size 1x1x1mm) was acquired. Functional MRI data were recorded using a gradient echo planar imaging sequence (echo time = 30ms, repetition time (TR) = 2s, flip angle = 75°, field of view = 220 x 220mm, matrix = 72x72, 32 slices, acquisition order interleaved odd first, slice thickness = 3mm, and 1mm interslice gap; 350 dynamic scans resulting in 700s acquisition time).

We performed image preprocessing according to recent publications ^17,19^. We removed the first five volumes to ensure that steady-state magnetization had been reached, and performed slice-time correction, realignment, and co-registration of functional images to T1-weighted data. Tissue class segmentation of the T1-weighted data for grey and white matter was performed using SPM12 with default parameters (htttp://www.fil.ion.ucl.ac.uk/spm) for creating individual subject grey matter masks (see below). Finally, we removed quadratic temporal trends in a voxel-wise fashion from the fMRI data using the Data Processing Assistant for Resting-State fMRI (DPARSF, http://rfmri.org/DPARSF) ^50^.

### Signals and Outcomes

#### Global grey matter fMRI signal

Functional MRI signals were extracted from subject-specific masks. For extracting the gGM fMRI signal, GM probability maps were registered to fMRI data using DARTEL (Diffeomorphic Anatomical Registration Based on Exponentiated Lie Algebra, neurometrika.org/) ^51^, available in SPM12 and thresholded at 50% GM probability. The negative temporal derivative (-d/dt) of the extracted gGM fMRI signal was generated by computing its first derivative and multiplying it by -1. All negative values were set to zero following methods described by ^17^ (**Fig. S5**). The average slope of the rising/falling flank of the signal in Fig. S5G was calculated by averaging the negative (B→S CSF or S→B gGM fMRI) or positive (S→B CSF or B→S gGM fMRI) values of d/dt of the respective signal (gGM or CSF fMRI).

#### CSF signal

For extracting CSF signals, individual subject CSF masks were delineated in native space following the approach of Han and colleagues ^19^. To this end, voxels with the highest signal intensity were selected from the three bottom slices of the fMRI data. Anatomical accuracy of the masks was confirmed by comparison with the individual subjects’ T1-weighted structural data. Voxels containing the brainstem or cerebellum were excluded from the masks, resulting in a ring-like shape.

#### PFI signal

PFI signal masks were generated for each subject individually by creating an average image for all bursts, containing all time points starting at 5 TRs after burst onset identified by the EEG labels until the burst offset. A suppression image was generated by averaging all frames starting 5 TRs after the onset of all suppression epochs identified on the EEG labels until the end of the suppression phases, respectively (**Fig. S1a-c**). We used the images 10-5 TRs and 15-20 TRs after the respective EEG transition to analyze burst-suppression or suppression-burst transitions. The suppression image was subtracted from the burst image, and the result (**Fig. S1d**) was binarized using a z-score threshold of -1.5 (**Fig. S1e**). The resulting mask was intersected with a freesurfer-generated mask of the ventricles, which had been centrally eroded to remove intraventricular voxels (**Fig. S1f,g**).

For each subject, we extracted their voxel-average gGM, PFI and CSF fMRI time course signals across their masks from the preprocessed fMRI data. The extracted time courses were temporally filtered at 0.1 Hz and smoothed using a moving average over five timepoints for visual presentation. For statistical analyses and comparison between subjects, time courses were intensity normalized (z-scored).

### Dataset 2: Hypercapnic challenge experiment

#### Participants

The experiment was performed in 21 healthy participants (age 29.1±8.6y, 10 female). Data from 17 subjects were included in these analyses; 4 participants were excluded due to corrupted data (n=1) or movement artifacts (n=3).

#### Experimental conditions and event definition

For the CO2 challenge, medical air (21% O2, 79% N2) or hypercapnia (medical air + 5% CO2) was applied using a sealed face mask and a gas mixer (Altitrainer, SMTec, Switzerland). A gas analyzer (ML206, AD Instruments, USA) facilitated end-tidal CO2 and O2 measurements. Hypercapnia was applied in 15 min runs with alternating blocks of 3 min air and 3 min hypercapnia (5 % CO2) (**Fig. 2A**). Because of a delayed delivery (about 5 m tubing) and ramping times of the gas mixer, it took about 30 sec ramp time until steady-state CO2 levels were reached.

#### MRI acquisition

Data were acquired on a 3T Philips Ingenia Elition X using a 32-channel head coil. T1-weighted MPRAGE (TE = 4 ms, TR = 9 ms, α = 8°, TI = 1000 ms, shot interval = 2300 ms, SENSE AP/RL 1.5/2.0, 170 slices, matrix size = 240x238, voxel size = 1x1x1 mm³) data were used for tissue type segmentation and generation of GM masks.

Pseudo-continuous ASL (pCASL) MRI was acquired for 15 min. Sequence parameters were set following recent recommendations by the ISMRM perfusion study group ^52^ using a label duration (LD) of 1800 ms and a post-label delay (PLD) of 1800 ms. Image readout employed 2D gradient echo EPI (TE = 11ms, TR = 4500s, 20 slices, SENSE factor = 2, acquired voxel size 3.3x3.5x6 mm^3^, 100 dynamics, 15:09 min). Additionally, proton density-weighted (PDw) M0 data were acquired for quantification of cerebral blood flow (CBF) in ml/100g/min. BOLD-fMRI data were acquired in the same subjects in a separate run using a T2*-weighted gradient echo EPI sequence (TE = 30ms, TR = 1200s, 40 slices, SENSE factor = 1.5, multi-band = 2 (slice order: ascending foot to head), EPI-factor = 20, acquired voxel size 3x3x3 mm^3^, slice gap = 0.2 mm, flip angle = 70°, 750 dynamic scans, 15:04 min).

#### pCASL pre-processing

For image registration and spatial transformations, CBF time series were derived using custom MATLAB algorithms and SPM12 (Wellcome Trust Centre for Neuroimaging, UCL, London, UK). pCASL label and control images were motion-corrected using rigid body transformation routines. Label and control images were subtracted, and intensity values were normalized by M0 maps. Time series of subtracted signals was then quantified for CBF ^52^and a Gaussian filter with a 6 mm FWHM kernel size applied.

### Signals and Outcomes

T1-weighted data were used to generate grey matter masks as described above. DARTEL affine registration was not used for this dataset. GM-average CBF time courses were extracted per subject.

#### CBF

To calculate group average CBF maps per condition or block-events (i.e., normocapnia, hypercapnia, N→H or H→N transitions) as displayed in Fig.2 C, we averaged spatially normalized CBF maps from time points showing minimum (for normocapnia) and maximum (for hypercapnia) values of each condition across subjects excluding the first ∼30 seconds at the beginning and at the end of the segment to allow CO2 concentrations to stabilize. Analyses of CBF differences between normal- and hypercapnia are based on the average CBF values extracted from these selected periods across all subjects.

#### fMRI

gGM and CSF fMRI signal extraction was performed as described above for the burst-suppression data set. The realignment step for this data was done only by estimating the movement parameters but avoiding unwarping procedures. This was done to prevent modifications in the bottom slice using the SPM realignment option Estimate & Unwarp‘.

### Statistics

We used two-tailed t-tests, ANOVAs, or a Kruskal-Wallis test to compare statistical groups. The corresponding test is indicated in the respective figure legends. Normal distribution was tested with a Shapiro-Wilk Test. The significance threshold was set to p < 0.05. To investigate analysis. In the hypercapnic challenge in **Fig. 2e**, we calculated the Pearson’s correlation between the gGM and CSF fMRI signals from 10 s (H→N) or 30 s (N→H) after switching the gas mixer for 48 sec. To investigate the correlations between the gGM, PFI and CSF fMRI signals in **Fig. 1h and i**, we used a bivariate Spearman correlation analysis. In the slice-sensitive CSF signal analysis, we performed repeated-measures ANOVA per event for each time point separately during the 30s following the respective transition.

## Funding

German Research Foundation (395030489) - CS and CP German Research Foundation (DFG SFB/TRR167 B07) - JP.

BZ is an Albrecht-Struppler-Clinician Scientist Fellow, funded by the Federal Ministry of Education and Research (BMBF) and the Free State of Bavaria under the Excellence Strategy of the Federal Government and the Länder, as well as by the Technical University of Munich - Institute for Advanced Study.

## Author contributions

Conceptualization: JZ, CS, BZ

Investigation sevoflurane dataset: JZ, CB, CE, SS, VN, MB, GS, AR, RI, DG, RN, DMH, JP, CS, BZ

Investigation hypercapnia dataset: JZ, CB, CE, GH, RN, LS, JK, SS, LS, JK, SK, CZ, CP, CS, BZ

Visualization: JZ, CB, CE, GH, VN, MB, CS, BZ

Funding acquisition: CZ, CP, CS, BZ Supervision: CP, CS, BZ

Writing – original draft: JZ, CB, CE, CP, CS, BZ

Writing – review & editing: all authors

## Competing interests

Authors declare that they have no competing interests.

## Data and materials availability

All data are available in the main text or the supplementary materials. Raw data will be made available by the corresponding authors upon reasonable request.

## Notes

### Competing Interest Statement

The authors have declared no competing interest.

### Summary of Updates

In the revised version, we included a novel technique we developed to detect CBV-associated changes of total brain volume. It is based on detecting partial volume effects at the border of the lateral ventricles (parenchyma-fluid interface, PFI), which gets shifted at burst-suppression transitions. The title, introduction, methods, discussion, and figures were revised and updated accordingly. Supplemental Materials were updated as well.

