## Supplementary materials for "Total cerebral blood volume changes drive macroscopic cerebrospinal fluid flux in humans"

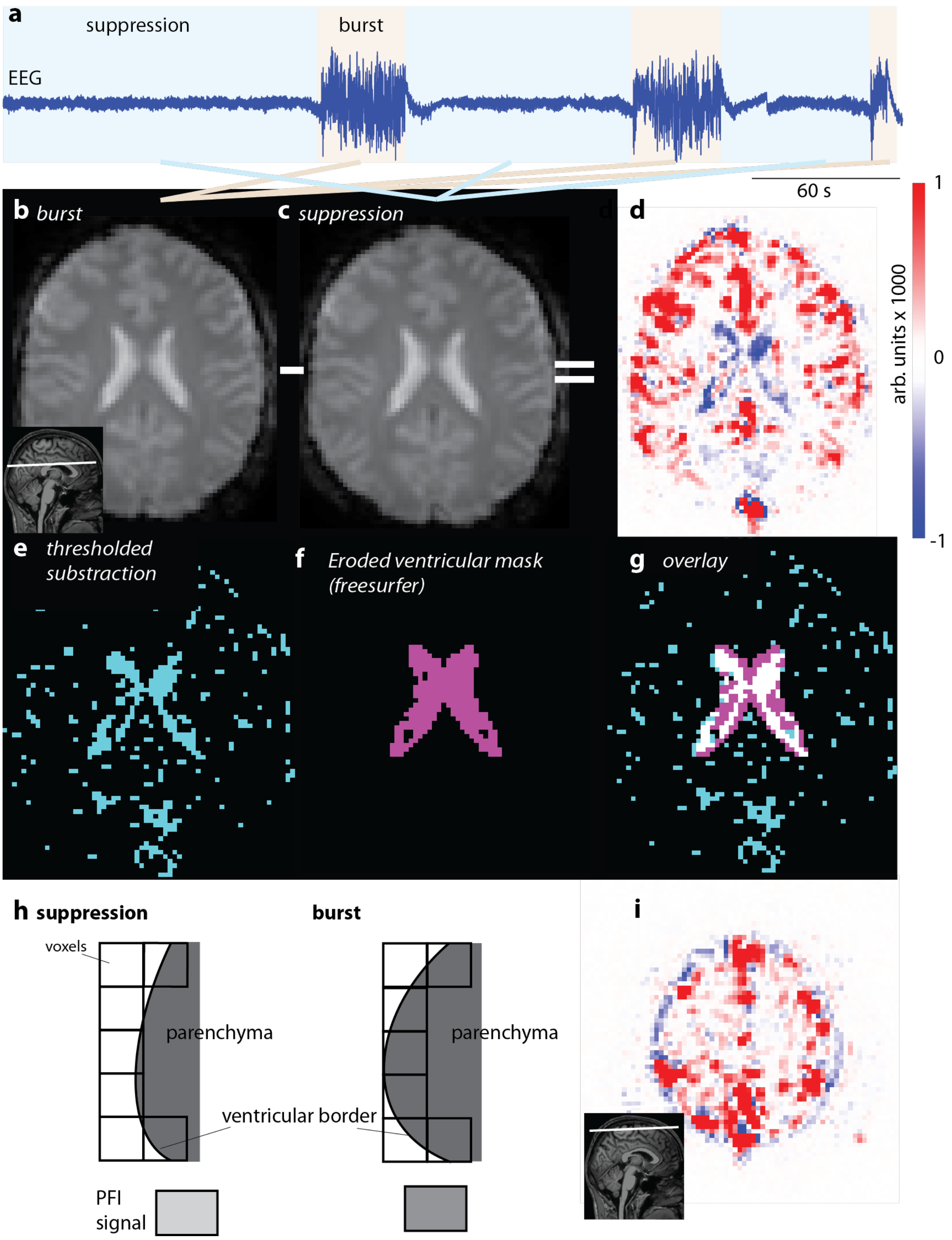


**Fig. S1: Generation of the parenchyma-fluid-interface (PFI) mask** (**a**) EEG trace from a representative subject (#5) with shaded suppression (*blue*) and burst (*orange*) epochs. (**b-c**) averages of all EPI image frames from burst (b) and suppression (c) epochs. The inset in (b) depicts the slice position. (**d**) Normalized subtraction image of the images in (b) and (c). Red voxels indicate higher signal intensity during bursts and are located in the grey matter and large blood vessels, while blue voxels have higher signal intensities under suppression and are located mainly at the border of the lateral ventricles. (**e**) Thresholded (< 0) and binarized subtraction image (d). (**f**) Freesurfer-generated ventricular border mask. (**g**) Overlay of the masks in (e) and (f) reveals very high agreement (*white voxels*). (**h**) Schematic depiction of the partial volume effects underlying the PFI contrast. Although the spatial resolution of the EPI scan is not high enough to resolve the movement of the ventricular border in detail, partial volume effects cause intensity differences between burst- and suppression-states. (**i**) Subtraction image as in (d), at a more cranial slice position (*inset*). Note the ring of blue (negative) voxels surrounding the positive cortical voxels.

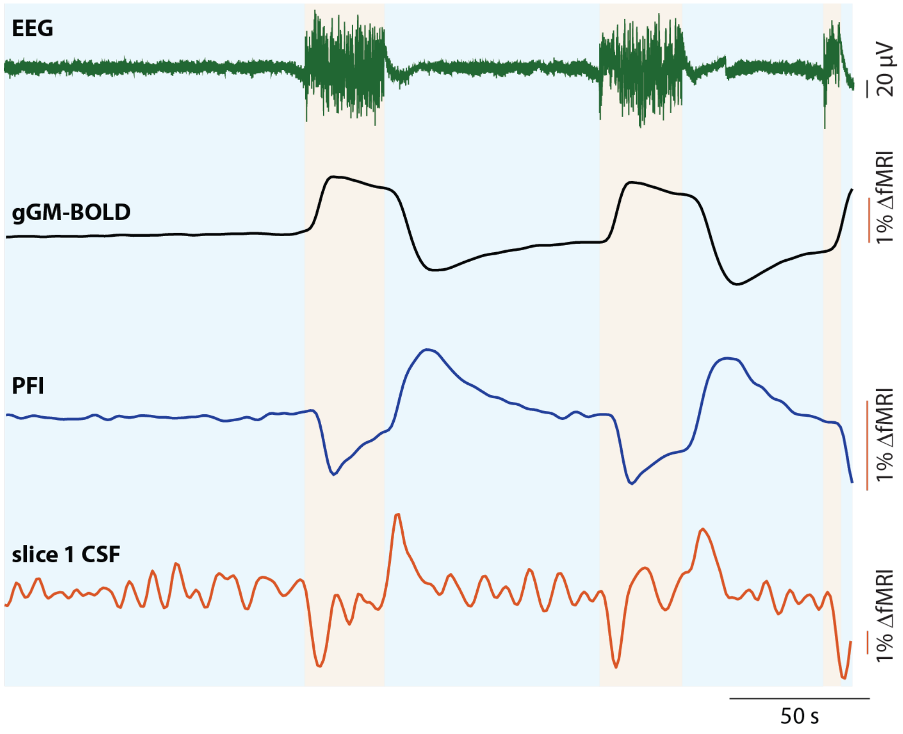


**Fig. S2: Co-registered EEG and fMRI signals from an additional subject (# 5).** Simultaneously recorded EEG (*green*), gGM (*black*), PFI (*blue*) and slice 1 CSF (*red*) fMRI signals. The suppression (*light blue*) and burst (*orange*) epochs are shaded in the background.

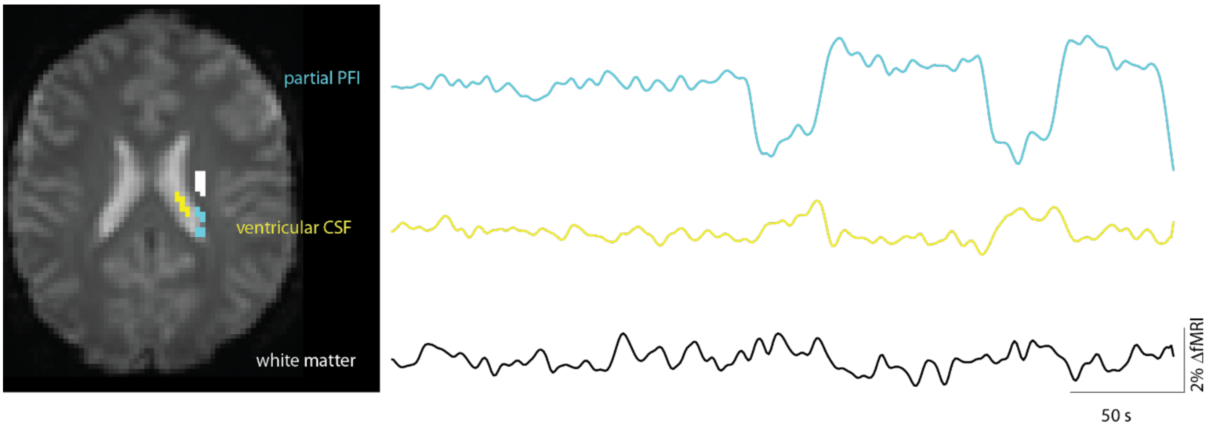


**Fig. S3: The PFI signal is not influenced by signal from the surrounding white matter or the ventricular CSF**. left: representative EPI-slice from subject #5. Overlayed are representative masks containing similar voxel numbers in PFI (*blue*), ventricular CSF (*yellow*) and white matter (*white*). The corresponding intensity traces are shown on the right. Note that the PFI signal time course is neither reflected in the white matter nor in the ventricular CSF at the same slice.

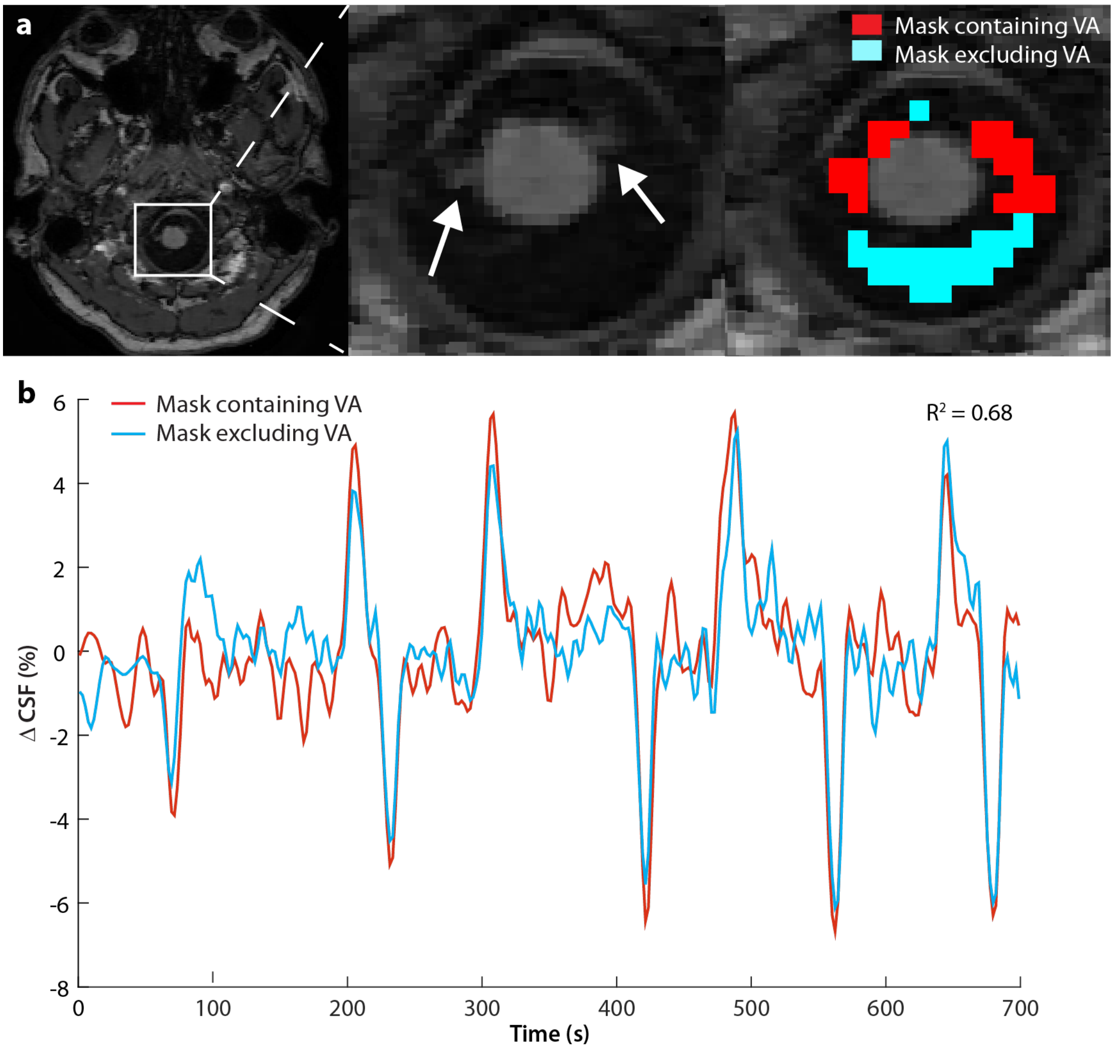


**Fig. S4: The CSF fMRI signal is not affected by influx of blood through the vertebral arteries.** (**a**) Representative axial T1-weighted image from subject #2 (slice 1, *left*) and enlargement (*middle*) of the CSF-containing region comprising the cisterna premedullaris, cisternae cerebellomedulares laterales and cisterna cerebellomedullaris posterior. White arrows point at the vertebral arteries. CSF masks (*right*) including (*red*) and excluding (*light blue*) the vertebral arteries. Note that the masks were adapted to contain the same number of voxels to obtain comparable signal-to-noise ratios. (**b**) CSF fMRI time courses extracted from the masks including (*red*) and excluding (*light blue*) the vertebral arteries.

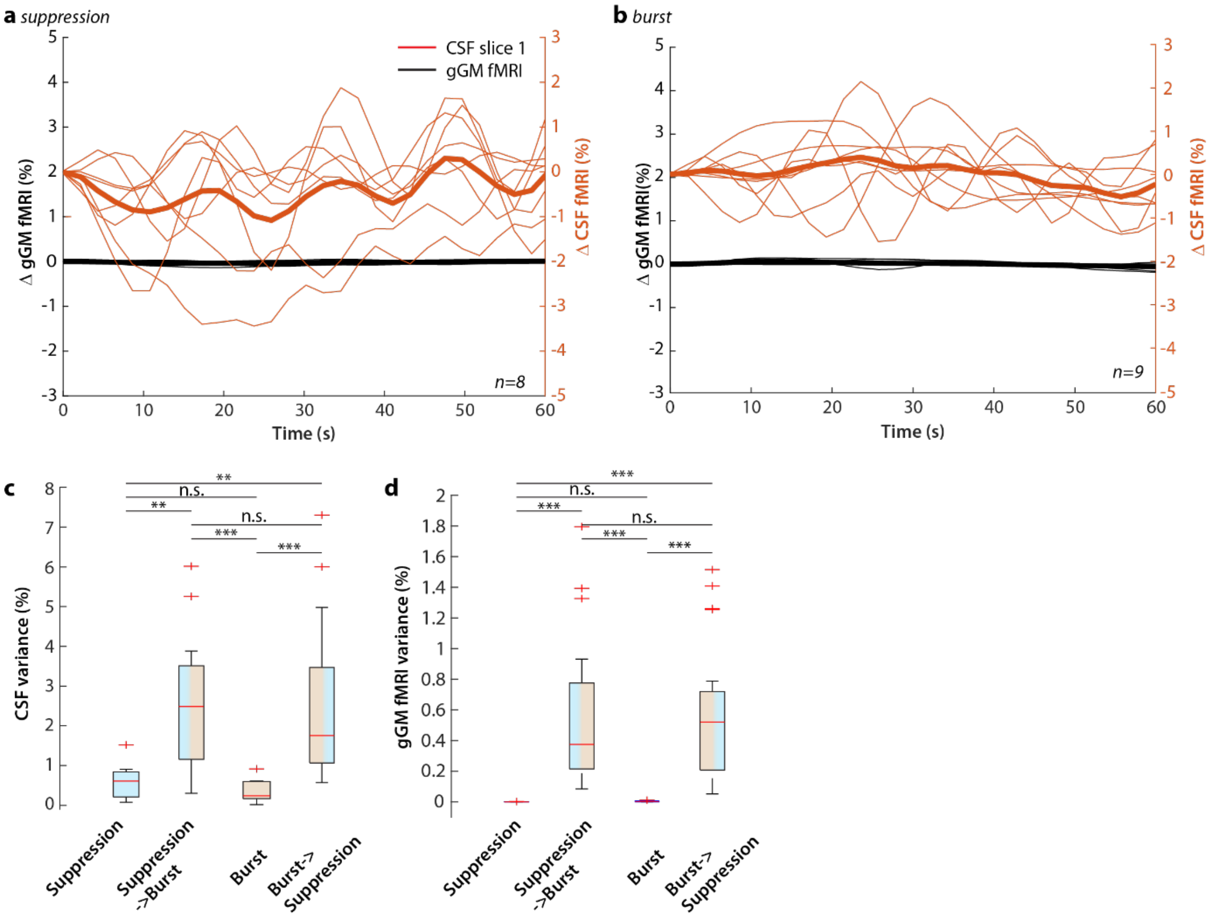


**Fig. S5. Global-GM and CSF fMRI signals during steady-state conditions. (a)** Time courses of all steady state suppression periods: gGM (*black*) and CSF (*red*) fMRI signals. Thick lines represent the average, fine lines the individual events (n=8). (**b**) Same as (a) for all steady state burst periods (n=9). (**c**) Box plots representing the variance of the CSF signal during suppression, S⟶B transitions, bursts, and B⟶S transitions. (**d**) same as (c) for the gGM fMRI signal. N.s. not significant. **p<0.005, ***p<0.001. Kruskal Wallis test with Dunn-Sidak post-hoc comparison.

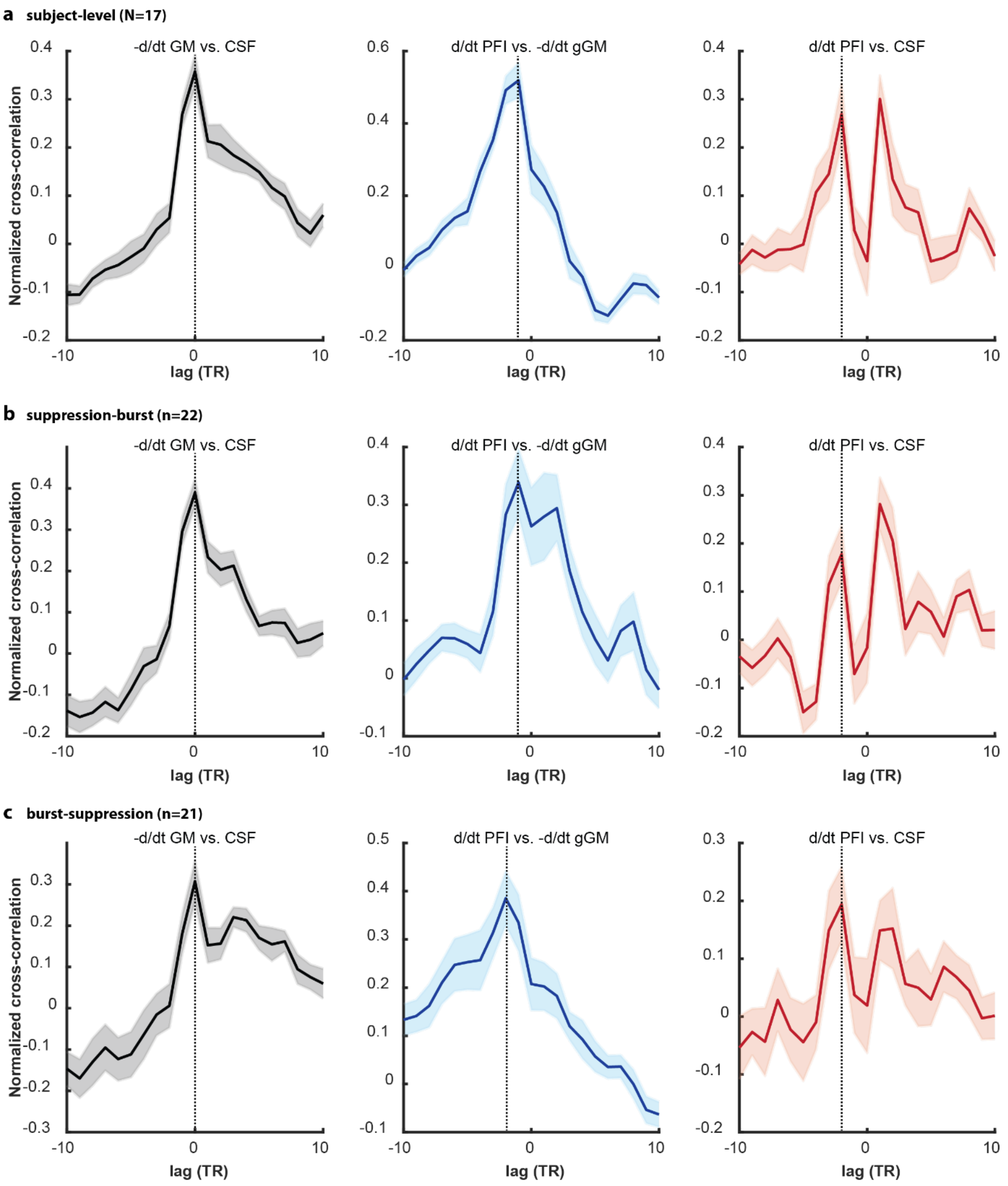


**Fig. S6: Cross-correlation analysis – the PFI signal precedes the gGM and CSF fMRI signals.** (**a**) Cross-correlation on a subject level (N = 17 subjects) for -d/dt GM vs CSF (*black, left*), PFI vs. -d/dt gGM (*blue, middle*) and d/dt PFI vs. CSF (*red, right*). The solid line depicts the mean, the shaded area represents SEM. Note that the second peak in the d/dt PFI cross-correlation is likely caused by the relatively large sinusoidal noise in the PFI signal. (**b**) Same as (a) for all suppression-burst transitions (n=22 events). (**c**) Same as (a) for burst-suppression transitions (n=21).

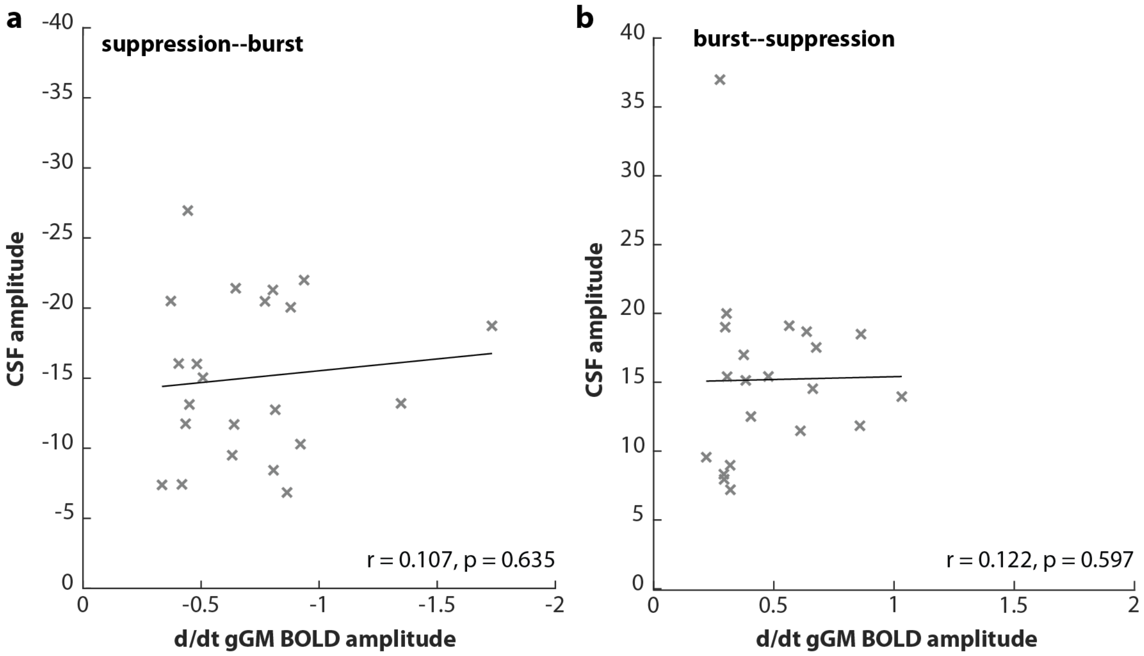


**Fig. S7: Control analysis for the specificity of PFI effect on CSF flow - No correlation between the d/dt gGM and the CSF amplitude**. (**a**) Scatterplot of the d/dt gGM and the CSF amplitude. Crosses represent individual transition events; the line depicts linear regression. (**b**) same as (a) for all burst-suppression events.

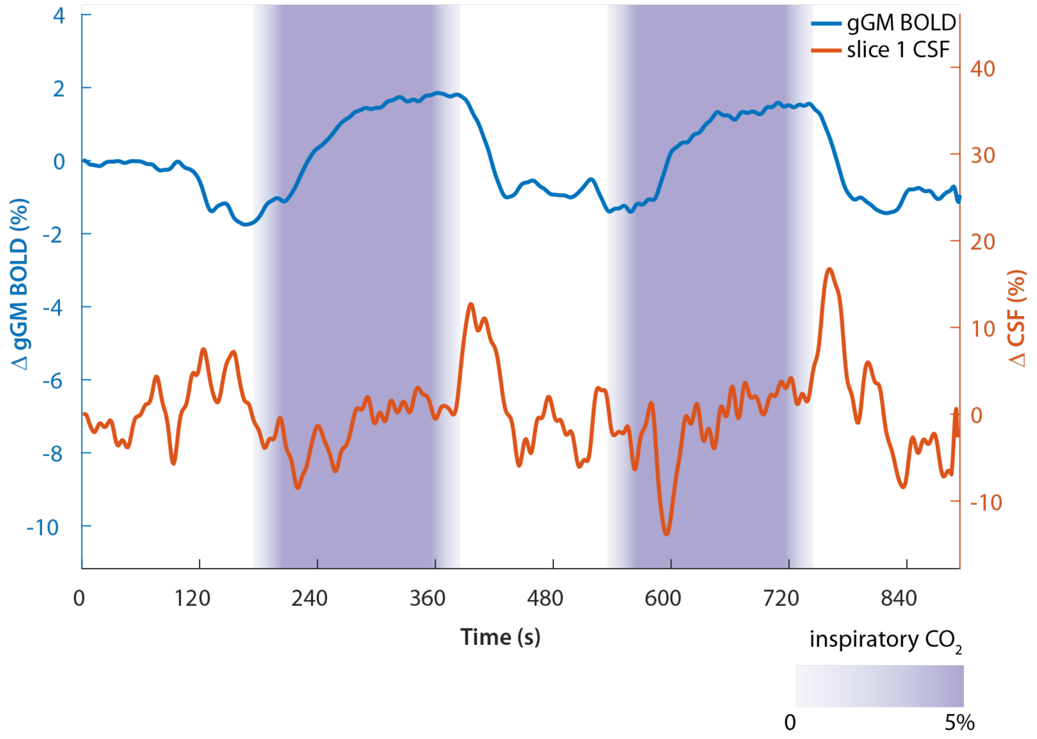


**Fig. S8: Coupled gGM and CSF fMRI signal time courses induced by a transient hypercapnic challenge – data from a representative subject.** Representative gGM *(top, blue)* and CSF fMRI signal *(bottom, red)* time courses. $CO_2$ application periods are indicated in purple.

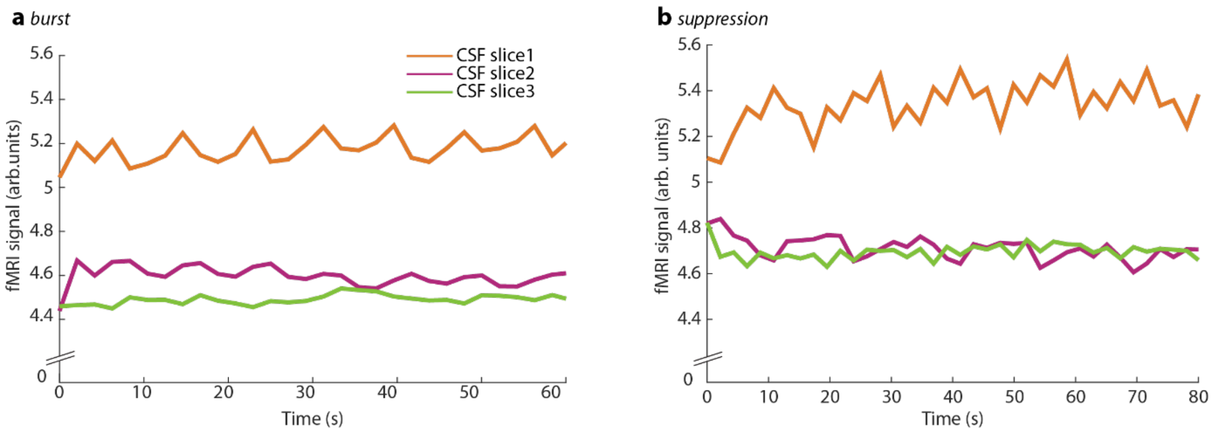


**Fig. S9. The steady-state CSF signal is higher in slice 1 than in slices 2 and 3.** (**a**) Representative CSF signal during a state burst period. Slice 1 (*orange*), slice 2 (*pink*), slice 3 (*green*). (**b**) Same as (a) for a representative steady state suppression period.

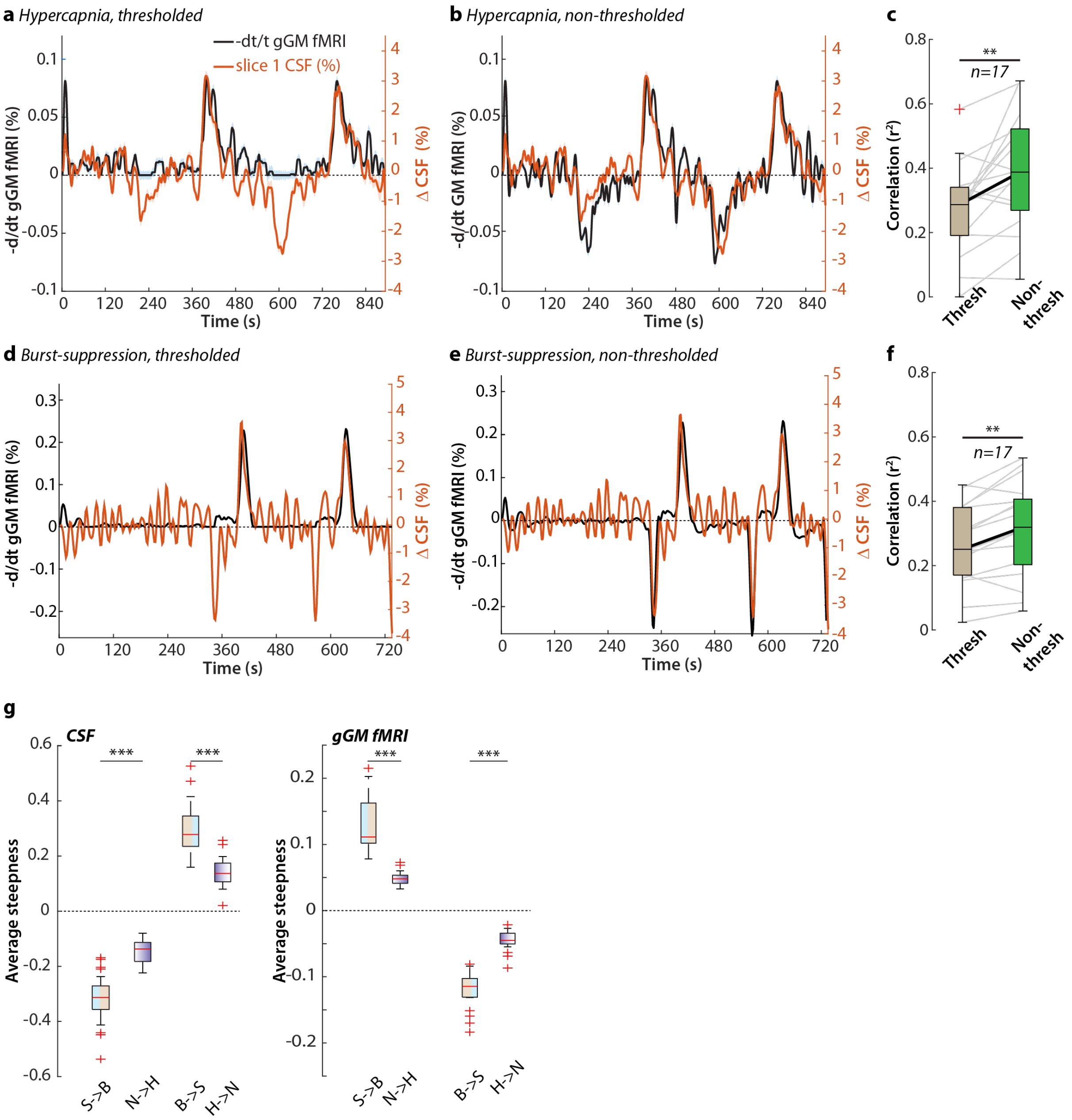


**Fig. S10: Relation of the gGM and CSF fMRI signals across experiments.** (**a**) Mean negative derivative (-d/dt) of the gGM fMRI signal, thresholded to zero for negative values (*black*) and mean CSF signal (*orange*) from all subjects *(n=17)* in the hypercapnia challenge experiment. (**b**) Same as (a) for the non-thresholded mean -d/dt gGM fMRI signal. (**c**) Correlation between the CSF signal and the zero-thresholded -d/dt gGM signal (left, brown) as well as between the CSF signal and the non-thresholded -d/dt gGM signal (right, green) for all subjects (n=17) in the hypercapnia challenge experiment. Box plot and individual values (grey lines). (**d**) -d/dt gGM fMRI signal thresholded to zero for negative values (*black*) and CSF (*orange*) time course of the signals from a representative subject (#5) in the burst-suppression experiment. (**e**) Same as (d) for the non-thresholded -d/dt gGM signal. (**f**) Same as (c) for all subjects in the burst-suppression experiment. **p<0.005. Paired t-test. (**g**) Averaged slope (d/dt) of the CSF signal in slice 1 (*left*) and the gGM fMRI signal (*right*) during the transition from suppression to burst (n=22), normo- to hypercapnia (n=17), burst to suppression (n=21) and hyper- to normocapnia (n=17). ***$p<0.001$. **$p<0.005$. Paired t-test (c, f), Wilcoxon rank sum test (g).
